# An adolescent neuroimaging database combining movie-watching, eye-tracking and cognitive tasks

**DOI:** 10.64898/2026.08.28.747840

**Authors:** J. Bužinel, J.Y.S. Choy, J. Norman, J. Hughes-Nind, E. Levchenko, F. Dick, J.I. Skipper, C.O. Carlisi

**Author notes:** These authors contributed equally to this manuscript (J. Bužinel, J.Y.S. Choy).

## Abstract

Adolescence is a critical period of neurodevelopment, yet most neuroimaging datasets focus on adult populations, leaving a gap in our understanding of how the brain processes information during this formative stage. Here we present a multimodal neuroimaging dataset acquired from 41 adolescent participants aged 11–18 years, combining 3T fMRI data with concurrent eye-tracking and physiological monitoring during naturalistic movie-watching. Data are shared in BIDS-compliant format and technical validation demonstrates good data quality across participants, with low head motion and strong inter-subject neural synchronisation during movie-watching. In addition to the scanning session, participants completed remote assessments covering a broad range of self-reported developmental and mental health traits, alongside cognitive tasks targeting reward-based and social learning. This dataset offers a rich resource for studying the adolescent brain, with particular utility for research on individual differences in mental health and cognition. All data and processing code is openly available to facilitate reproducible science.

## Background and Summary

Adolescence is a sensitive developmental period characterised by ongoing maturation of prefrontal and limbic brain circuitry underlying emotion regulation, reward processing, and social behaviour (Blakemore, 2008; Casey et al., 2008; Paus et al., 2008). It is also the period during which many mental health problems and psychiatric disorders first emerge, with the majority of lifetime cases across all mental disorders beginning before age 25 (Kessler et al., 2005; Solmi et al., 2022) and thought to be due in part to the ongoing maturation of key neural systems through adolescence (Galván, 2017.). Understanding how the brain develops during this period is therefore critical for characterising developmental trajectories and identifying factors that may contribute to mental health risk.

Previous large-scale developmental cohorts such as the Adolescent Brain Cognitive Development Study (ABCD; Casey et al., 2018), Lifespan Human Connectome Project Development (HCP-D; Somerville et al., 2018), and Healthy Brain Network (HBN; Alexander et al., 2017) have transformed the field of developmental neuroscience by providing structural, diffusion, resting-state, and task-based MRI from thousands of young people alongside extensive phenotyping. However, these datasets rely predominantly on restingstate acquisitions and short, tightly controlled experimental tasks. Naturalistic paradigms (e.g., movie-watching) provide a complementary approach for studying brain function as participants instead engage with dynamic stimuli that unfold continuously over time. Compared with task-based fMRI, movie-watching has been shown to better predict individual differences in cognition and affect from functional connectivity patterns (Finn and Bandettini, 2021). They are also particularly well-suited to developmental neuroimaging because they sustain attention, improve engagement, and reduce in-scanner head motion in children and adolescents (Vanderwal et al., 2015, 2019), while providing greater ecological validity and evoking reliable, temporally synchronised neural responses across individuals (Hasson et al., 2004).

Existing naturalistic developmental datasets have, however, typically used short movie clips, which can capture responses to individual events but provide limited opportunity to examine how neural responses develop across longer, continuous experiences. Publicly available developmental datasets using full-length movies remain relatively uncommon, despite their potential to capture events, characters, and social interactions as they occur over time in ways that more closely resemble everyday life. Indeed, such paradigms have been shown to be especially informative for studying processes central to adolescent development, including theory of mind and social-cognitive network maturation (Richardson et al., 2018), emotion processing, and the development of large-scale functional networks (Moraczewski et al., 2020).

To address these limitations, a small number of publicly available datasets have begun to combine naturalistic stimuli with behavioural and cognitive assessments. For example, *StudyForrest* (Hanke et al., 2016) provides 3T fMRI data alongside eye-tracking and physiological recordings of participants watching the movie *Forrest Gump*, while the *Grand Budapest Hotel* dataset focuses on social cognition (Visconti di Oleggio Castello et al., 2020). Notably, the *Naturalistic Neuroimaging Database* (NNDb v1.0; Aliko et al., 2020) provides movie fMRI data from 86 adults, each watching one of ten feature films, accompanied by a battery of behavioural testing. Its extension, NNDb-3T+ (Levchenko et al., 2026), expands on this by providing rich multimodal data from 40 adults watching *Back to the Future* with concurrent physiological and eye-tracking recordings, and a cognitive battery. To date, however, such resources have focused predominantly on adults. Adolescents differ from adults not only in brain structure and function, but also in developmental characteristics highly relevant to mental health research, including pubertal maturation, peer-sensitive social learning, reward sensitivity, and rapidly evolving emotion regulation capacities.

Here, we address this gap by presenting a multimodal naturalistic neuroimaging dataset acquired from 41 adolescents aged 11–18 years. Extending the naturalistic movie-watching approach of NNDb v1.0 (Aliko et al., 2020) and NNDb-3T+ (Levchenko et al., 2026) to adolescence, this dataset, the *Naturalistic Neuroimaging Database Teens* (NNDb-Teens) is, to our knowledge, the first publicly available dataset in adolescents of its kind to combine full-length movie-watching (*Back to the Future*, with 3T fMRI, 2×2×2mm resolution) with concurrent eye-tracking and physiological monitoring, behavioural assessments (questionnaires on mental health symptoms, well-being, emotion regulation, pubertal development), cognitive tasks (reward-based and social learning), and experience sampling methods. We expect that the dataset will support an array of future studies on individual differences in naturalistic brain function and, in conjunction with the NNDb-3T+, on the cross-sectional developmental differences in naturalistic processing between adults and adolescents. All raw and preprocessed imaging data, physiological recordings, quality-control metrics, and processing code are openly shared in BIDS-compliant format to facilitate reproducible developmental neuroscience research.

## Methods

### Participants

41 participants (*mean*(*SD*)*_age_* = 15.10(1.96)*years, range_age_* = 11 *−* 18 *years, n _f emale_* = 31) completing the fMRI scan represent a subset of participants from a larger study (preregistered at https://osf.io/uaf63). Participants were recruited primarily through local secondary schools across England and Wales, but also community organisations, local libraries and word-of-mouth referrals. In some cases, schools were contacted directly by researchers, and in others, schools contacted the research team after seeing an advert on the lab website. Information sheets about the study was also sent to school staff, who distributed them among the students and their parents/carers. For participants who were not recruited via a school, the information sheets covering the same content were shared via email with the child and parent. Interested children (and their parents if the child was under 16) then completed an online consent form via a secure and encrypted website, where they indicated whether they would be interested in taking part in the fMRI part of the study and completed a separate consent form for the fMRI scanning.

Participants were prescreened for MRI safety eligibility and reported normal or correctedto-normal vision and hearing. The sample comprised native English speakers aged between 11 and 18 years with no history of neurological disorders.

The full study protocol, including MRI data acquisition, was approved by the University College London Research Ethics Committee (Ethics Project ID number: 22135/003). Upon completion, participants received £45 in voucher compensation.

### Procedure

Prior to their visit, participants completed an MRI safety screening form assessing the presence of metal implants, pacemakers, and other potential contraindications for MRI scanning. Participants also completed behavioural data collection over a period of two one-week windows, six to 12 weeks apart. During the first week, participants completed an online battery of 12 questionnaires collecting demographic and mental health information, as well as two cognitive tasks delivered through Gorilla (https://gorilla.sc) and Pavlovia (https://pavlovia.org). Experience Sampling Methodology (ESM) surveys (five per day for seven days) were completed twice, once during week one and again in week 2, approximately six to twelve weeks later. The behavioural data collection procedure is described in more detail on OSF (https://osf.io/uaf63).

FMRI scans were scheduled either between the two data collection windows or after the second week. Upon entering the scanning room, participants selected appropriate sized earbuds for use with noise-attenuating headphones and wore a hairnet to prevent hair from interfering with the latches of the 30-channel head coil. Participants then inserted the earbuds, laid supine on the scanner bed, and positioned their head within the coil. A pillow was placed beneath the legs to enhance comfort and minimise movement. Head motion was further restricted using an in-house developed stabilisation helmet with inflatable cushions (MR-MinMo; patent number GB 2205139.5, filed 07 April 2022).

The participant’s head position was localised, and a front-surface mirror was mounted on the radiofrequency head coil to facilitate visual presentation of the movie. Once positioned inside the scanner bore, the lights in both the bore and scanning room were turned off. A short movie segment was presented to confirm adequate visual and auditory perception, and audio volume was individually adjusted. A brief localiser scan was subsequently acquired, and the field of view was adjusted to ensure whole-brain coverage; when full coverage was not achievable, the operator excluded the minimum number of cerebellar slices necessary, consistent with the protocol used in NNDb-3T+ (Levchenko et al., 2026).

Following setup, the movie presentation script was initiated and eye-tracker calibration and validation procedures were completed. The operator adjusted parameters as needed to achieve optimal validation quality before commencing the movie-viewing. Participants viewed the full film *Back to the Future* (Zemeckis, 1985), presented in three segments of similar length. Eye-tracker calibration was repeated prior to each segment. During breaks between segments, participants could request to exit the scanner; however, operators encouraged participants to remain positioned to minimise head displacement whenever possible. After the scan, participants completed a short survey about their engagement with the movie, different aspects of mood during the movie, and whether they had previously watched the movie. The complete procedure lasted approximately three hours. An overview of the study procedure, preprocessing, and validation steps is provided in Figure 1. Data completeness for each participant across the questionnaire, ESM, cognitive task, and movie-watching components is also summarised in Supplementary Table S1.

**Figure 1:**
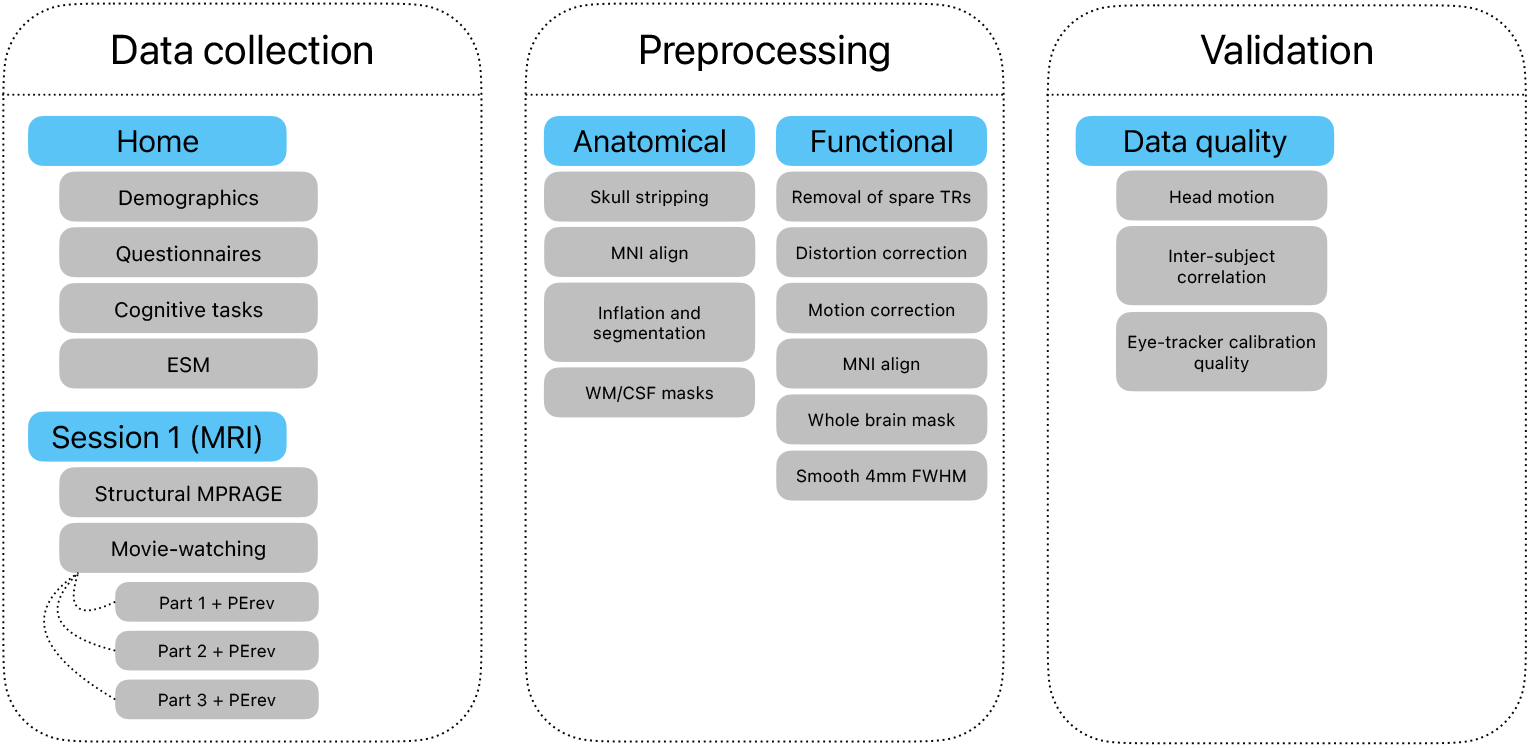
Overview of the study. 1) Data collection: participants completed a battery of questionnaires and cognitive tasks at home. The fMRI data were acquired during a single session, including a 5-minute structural MPRAGE scan and a movie-watching task split into three functional runs (parts). 2) Preprocessing: the structural and functional epi (movie-watching) data were preprocessed as outlined in the figure. 3) Validation: data quality and validity were assessed by looking at head motion, inter-subject correlation during movie-watching, eye-tracker calibration quality for each run of the movie.

### Questionnaires

The questionnaire battery required approximately 30 minutes to complete and was administered via smartphone app to be completed in participants’ own time, prior to coming in for their scan. The questionnaires collected information on basic demographics, mental health, well-being, emotion regulation, and quality of life.

The first section collected information on basic demographics. The second section comprised eleven validated psychological questionnaires selected to provide a comprehensive assessment of mental health and well-being. All selected measures are well validated, reliable, and efficient, enabling broad coverage of psychological functioning whilst minimising participant burden. Questionnaires were administered via the eMoodie app (https://emoodie.com) and the Gorilla platform and are described below.

The *Patient Health Questionnaire* (PHQ-8) is an eight-item screening instrument widely used to assess depressive symptoms and related mental health conditions in primary care and research settings (Kroenke et al., 2001, 2009). Items are rated on a four-point scale ranging from 0 (“not at all”) to 3 (“nearly every day”). Despite its brevity, the PHQ-8 demonstrates sensitivity and specificity comparable to longer depression assessments. In this study, an 8-item version of this questionnaire was used, omitting the suicidality item for safeguarding reasons. The PHQ-8 has good psychometric properties for identifying depression and measuring depressive symptoms.

The *Generalized Anxiety Disorder scale* (GAD-7) is a seven-item measure used to screen for and assess the severity of anxiety symptoms in clinical and research contexts (Spitzer et al., 2006). Each item is rated on a scale from 0 (“not at all”) to 3 (“nearly every day”). Generalized anxiety disorder is among the most prevalent anxiety conditions in both clinical practice and the general population.

The *Strengths and Difficulties Questionnaire* (SDQ) is a 25-item measure comprised of five subscales: 1) emotional symptoms, 2) conduct problems, 3) hyperactivity/inattention, 4) peer relationship problems, 5) prosocial behaviour (Goodman, 1997). Scales one through four can be added together to generate a total difficulties score. The questionnaire is used to screen for child and adolescent mental health, and is particularly well suited to community samples with its emphasis on strengths.

The *Revised Child Anxiety and Depression Scale* (RCADS) is a 47-item measure comprised of six subscales covering: 1) separation anxiety disorder, 2) social phobia, 3) generalised anxiety disorder, 4) panic disorder, 5) obsessive compulsive disorder, and 6) low mood (Chorpita et al., 2000). The questionnaire also yields a total anxiety scale (sum of five anxiety subscales) and a total internalising scale (sum of all subscales). The questionnaire shows good reliability on subscales and total scale.

The *Difficulties in Emotion Regulation Scale* (DERS) is a 36-item measure comprised of six subscales (non-acceptance of emotional responses, difficulties engaging in goal-directed behaviour, impulse control difficulties, lack of emotional awareness, limited access to emotion regulation strategies, and lack of emotional clarity) used to assess how respondents relate to their emotions (Gratz and Roemer, 2004). Items are rated on a 5-point Likert scale from ‘Almost never’ to ‘Almost always’.

The *Perceived Stress Scale* (PSS) is a 10-item measure exploring how different situations affect respondents’ feelings and perceived stress (Cohen et al., 1983). Participants rate items about their experience of stress (e.g., ‘… been upset because of something that happened unexpectedly’) in the last month on a 5-point Likert scale from ‘Never’ to ‘Very often’.

The *Reflective Functioning Questionnaire for Youth* (RFQ) is a 6-item measure evaluating the capacity to imagine and recognise one’s own and others’ mental states (Sharp et al., 2022). Problems with reflective functioning are involved in various forms of psychopathology and behavioural problems (e.g., Batenburg et al., 2026). The dataset includes the six-item version, however, users may wish to use a five-item version which has been found to perform better in adolescents (RFQ-Y-5; Sharp et al., 2022) and can be obtained by removing the first of the six items.

The *KIDSCREEN-27* is a 27-item measure comprised of five subscales: 1) physical wellbeing, 2) psychological wellbeing, 3) autonomy and parent relations, 4) peers and social support, and 5) school environment (Ravens-Sieberer et al., 2007). Participants respond to items on a 5-point Likert scale ranging from ‘not at all’ to ‘extremely’, or ‘never’ to ‘always’ based on their last week. The questionnaire is psychometrically robust and shows good internal consistency (Robitail et al., 2007).

The *Family Affluence Scale-III* (FAS) is a 6-item measure designed to provide an estimate of a family’s socioeconomic status (C. E. Currie et al., 1997; Torsheim et al., 2016). Questions ask about, for example, the number of computers, cars, and bathrooms in a household. The responses to each of the items are then summed to obtain a total score ranging from 0 to 13. The questionnaire has been used extensively to examine the material status of children and adolescents and is an important predictor of health (C. Currie et al., 2024; Elgar et al., 2016).

The *CRAFFT* is a substance screening tool for adolescents. In the current dataset, data was collected for Part One, asking participants to report the number of days in the past year they have 1) drunk alcohol, 2) used marijuana in any form, or 3) used anything else to get high (Knight et al., 2002).

The *Pubertal Development Scale* (PDS) is an 8-item measure with males omitting three items and females omitting two (Koopman-Verhoeff et al., 2020; Petersen et al., 1988). Participants are asked to judge the progression of features of their pubertal development (e.g., growth of body hair) by answering whether it has not yet started, barely started, definitely started or seems complete. The PDS is reliable, provides a good indication of Tanner stages when categorised, and has been used extensively in research in adolescents.

### Experience Sampling Methodology

The ESM surveys were completed on the eMoodie (https://emoodie.com) application, in response to notifications received on their phone. The short surveys included 16-25 questions, depending on the answers given, and took about one minute to complete. The questions asked about (1) participants’ affect on a seven-point Likert scale, (2) experience of stressors (i.e., whether anything negative has happened in the last 30 minutes and how they felt about it), (3) experience of positive events (i.e., whether anything positive has happened in the last 30 minutes and how they felt about it), (4) what they were doing at the time of receiving the notification, and (5) the setting and social settings (i.e., where they were and who they were with). Participants received 5 prompts per day for a sampling period of 7 days. Sampling was repeated approximately 6-12 weeks after the first sampling period.

### Cognitive Tasks

Two cognitive tasks (two-armed bandit task and social evaluation learning task) were selected to assess threat/reward and social processing, reflecting key aspects of cognition relevant to adolescent mental health and providing the basis for examining individual differences in cognitive function. A detailed description of the tasks is available in the pre-registration for the project (https://osf.io/uaf63). Participants completed both tasks remotely and each task took about 20 minutes to complete.

#### Dragon’s cave

This task is a two-armed non-stationary bandit task and an adaptation of Aylward et al., 2019’s four-armed bandit task tailored to children and adolescents. In the task, participants are instructed to pick one of two caves (green or blue) for a dragon called ‘Rory’ to sleep in each night. Participants are told that Rory likes discovering gold (reward) and dislikes damp (punishment) and that they should therefore aim to obtain gold and avoid damp. On a given night (trial), each cave could contain gold, damp, both gold and damp, or neither. Participants are told that the likelihood of obtaining gold and damp fluctuated independently in each cave throughout the game. The position (left or right) of the caves on the screen is randomised so that each cave sometimes appears on the left and sometimes on the right. To provide further motivation, participants are shown the amount of gold collected on the screen, alongside a bar of Rory’s ‘firepower’, which reduces in size when damp is obtained. The bar is designed so that it is unlikely to ever fully deplete. Participants completed a total of 154 trials, including 4 attention checks and 150 task trials. After completing the task, participants were asked to indicate how motivated (0-100) they were to (1) collect gold, and (2) avoid damp. Response data collected includes trial-level choice of bandit (cave), side on which cave was presented (left or right), and the outcomes associated with the chosen cave (reward/punishment). Further information about the task and modelling procedures is available at https://osf.io/tyf38.

#### Social Evaluation Learning Task Revised (SELT-R; Lau et al., 2024)

This is a modified version of the SELT (Button et al., 2015). In the task, participants learn about how a virtual rater feels about them (self-referential condition) or an avatar (other-referential condition) through receiving positive and negative personality feedback (e.g., exciting, dull). There are four feedback contingencies (liked, liked-repeated, neutral, disliked), yielding a total of eight blocks (self-liked, self-liked-repeated, self-neutral, self-disliked, other-liked, other-liked-repeated, other-neutral, and other-disliked), each formed of 20 trials presented in random order. On each trial, participants are asked to provide a rating of how likely they think the rater would be to describe them (or an avatar) with a positive word on a scale from 0 to 100, with higher scores corresponding to a higher likelihood of the positive feedback (e.g., 80% likely to be caring, 20% likely to be uncaring). At the end of each block, participants are asked to provide a global rating of how much the rater liked or disliked them (or an avatar) in the block from 0 (completely disliked) to 100 (completely liked). Response data includes trial-level word choices, positivity ratings, feedback by the computerised rater, and global positivity ratings after each block.

### fMRI Session

Participants viewed the *film Back to the Future* (Zemeckis, 1985), which had a total duration of 1 hour, 51 minutes, and 16 seconds. Participants were instructed to remain as still as possible during scanning while watching the movie naturally.

The movie file was split into three runs using the ‘ffmpeg’ package:

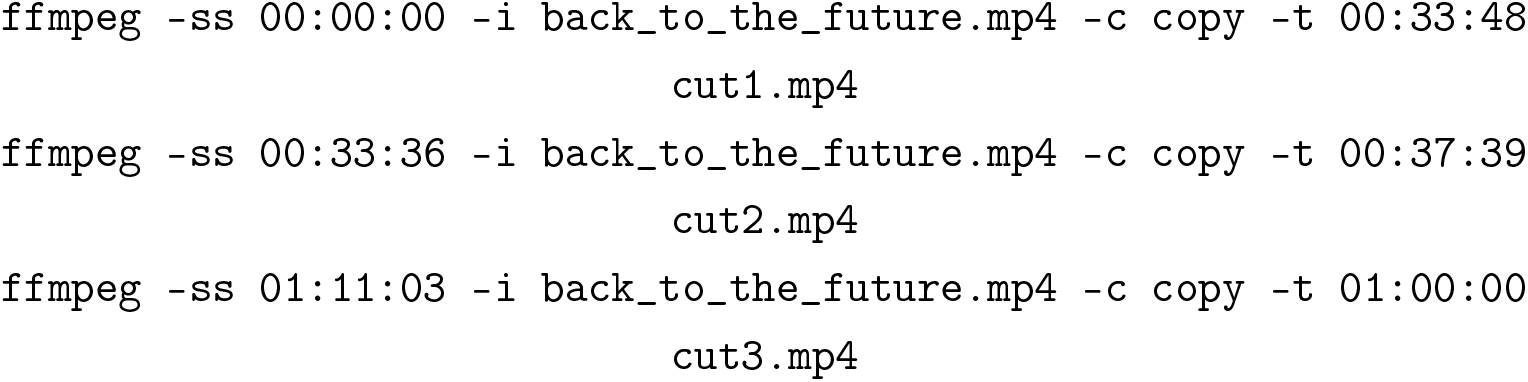

Cut points were selected to achieve comparable run durations while preserving smooth transitions between scenes. The second and third runs each included a 12-second overlap from the preceding segment to allow the hemodynamic response function (HRF) and associated psychological processes to theoretically return to a state comparable to that of the prior run. At the beginning of each run, eight initial TRs were acquired to allow stabilisation of the MR signal. Accordingly, 8, 16, and 16 TRs were removed from runs 1–3, respectively, during analysis so that the fMRI time series matched the duration of the full movie without temporal overlap. The final run lengths were 34 min, 38 min 03 s, and 40 min 12 s for runs 1–3, corresponding to 1360, 1522, and 1608 TRs, respectively.

The extracted video segments preserved the original audiovisual properties, retaining all frames without cropping or additional transformations. Video was encoded using H.264 (High profile), and audio was encoded in AAC (LC) format at a sampling rate of 48.0 kHz, bitrate of 339 kbps, and 5.1-channel output. The video resolution was 720 × 576 pixels with a 16:9 aspect ratio and a frame rate of 25 frames per second.

Movie presentation was controlled using a custom MATLAB script (MATLAB R2022b, version 9.13.0.204977) implemented with Psychtoolbox (v3.0.18) on a Windows PC (Windows 11 Pro v22H2, 64-bit) using GStreamer version 1.0.

### Data acquisition

Functional and anatomical MRI data were acquired for all sessions and tasks using a 3.0 T Siemens MAGNETOM Prisma scanner equipped with a 30-channel radio-frequency (RF) head coil (Siemens Healthcare, Erlangen, Germany).

#### MRI parameters: backtothefuture task

Functional images were acquired using a multiband echo-planar imaging (EPI) sequence with the following parameters: repetition time (TR) = 1500 ms, echo time (TE) = 35.2 ms, flip angle = 60°, field of view = 212 mm, voxel size = 2 mm isotropic, slice thickness = 2.0 mm, and 72 interleaved slices with an anterior–posterior phase-encoding direction (A→P). Echo spacing was 0.56 ms with a bandwidth of 2620 Hz/pixel, and a multiband acceleration factor of 4 was applied. The three movie runs comprised 1360, 1522, and 1608 TRs, respectively, including eight initial TRs acquired at the start of each run. A reverse phase-encoding scan (P→A) was acquired immediately following each run for distortion correction.

Following the functional scans, a high-resolution T1-weighted anatomical image was obtained using an MPRAGE sequence (TR = 2300 ms, TE = 2.98 ms, flip angle = 9°, field of view = 212 mm, voxel size = 1 mm isotropic, slice thickness = 1.0 mm, 208 sagittal slices, anterior–posterior phase encoding direction). Echo spacing was 7.1 ms with a bandwidth of 240 Hz/pixel. The anatomical scan lasted approximately five minutes.

#### Eye-tracker apparatus

Eye-tracking data were acquired using an MRI-compatible EyeLink 1000 Plus system with a long-range mount (SR Research Ltd., Mississauga, Ontario, Canada). The optical camera head (f = 50 mm, F1.4 lens) and infrared illuminator (FL-890) were positioned horizontally behind the projection screen inside the scanner bore on a dedicated mounting tray. A front-surface silvered mirror was attached to the anterior head coil to enable stimulus viewing. A binocular recording configuration was used with a sampling rate of 1000 Hz. Eye-tracker calibration employed a nine-point procedure covering the full display area. Accuracy validation required participants to fixate the same targets used during calibration (EyeLink software v5.15), and calibration–validation procedures were repeated until optimal accuracy was achieved.

Visual stimuli were presented in full-screen mode using a mirror-reversing LCD projector (EPSON LB-1100U) projecting onto a rear-projection screen, which participants viewed via the front-surface mirror mounted on the head coil. Participants were positioned 57.5 cm from the screen, which measured 35.5 cm × 26 cm. Stimuli were presented at native resolution and subtended a visual angle of 28.9° × 18.3°. The eye-tracker camera and illuminator were mounted behind the screen on an adjustable platform inside the bore. The positions of the screen, projector, eye-tracker platform, and mirror were standardised across participants and verified prior to each session.

Eye-tracking data were recorded during the *backtothefuture* task. Each experimental run began with eye-tracker calibration and validation initiated via a MATLAB script. Once calibration accuracy met predefined criteria, the experimental paradigm commenced following acquisition of eight initial TRs to allow signal stabilisation. The presentation script transmitted event markers (“MSG” entries in the EyeLink ASCII data file) indicating task onset. During the movie task, scanner pulses and video frame timings were logged to the eye-tracking data file (see Data Records for details). Stimulus presentation was implemented in MATLAB (R2022b; version 9.13.0.204977) using Psychtoolbox (v3.0.18) on a Windows PC (Windows 11 Pro v22H2, 64-bit) with GStreamer version 1.0.

#### Physiological recordings

Pulse oximetry data were recorded using a Siemens wireless peripheral pulse unit (Siemens Healthcare GmbH, Erlangen, Germany) during the *backtothefuture* task. The peripheral pulse unit was attached to the left index finger. Participants were instructed to avoid moving their hands and arms to prevent movement artefacts.

#### Post-session questionnaire

After completing the scan, participants completed a short questionnaire asking about their language background and their experience of the movie (e.g., how happy, excited, focused, bored they were).

### Preprocessing

Raw DICOM data were converted to Brain Imaging Data Structure (BIDS)-compliant format using the heudiconv tool (version 1.3.0; https://github.com/nipy/heudiconv) (Gorgolewski et al., 2016). All anatomical images were anonymised through facial feature removal using pydeface (version 2.0.2; https://github.com/poldracklab/pydeface). MRI data preprocessing was performed using the AFNI software suite (AFNI_23.0.03 “Commodus”) to ensure data quality and prepare the datasets for subsequent statistical analyses (Cox, 1996, Cox and Hyde, 1997).

### Anatomical

The anatomical T1-weighted image was skull-stripped and nonlinearly aligned to the MNI152 2009 template using the AFNI @SSwarper pipeline, yielding a standard-space anatomical image, a nonlinear warp field, and an affine transformation matrix. Cortical surface reconstruction was performed using FreeSurfer (recon-all, default parameters; version 7.3.2-20220804-6354275; http://www.freesurfer.net) (Destrieux et al., 2010; Fischl, 2012). The reconstructed surfaces were converted to AFNI-compatible format using the SUMA (Surface Mapper) tools and were subsequently used to generate white matter and ventricular regions of interest (ROIs) for use as nuisance regressors during preprocessing. These ROIs were eroded and incorporated as noise regressors in the preprocessing of the *Back to the Future* movie.

### Functional

Functional data underwent a series of preprocessing steps to improve alignment and reduce imaging artefacts. Initial volumes were removed from each functional run to eliminate pre–steady-state signal effects. The first eight TRs were removed from run 1 and the first sixteen TRs from runs 2 and 3.

Geometric distortions arising from susceptibility-induced field inhomogeneities were corrected using a blip-up/blip-down approach with reverse phase-encoding field maps. Median volumes from the forward and reverse phase-encoding scans were extracted, and midpoint warp fields were estimated and applied to the functional data using 3dNwarpApply. Motion correction was performed using a hierarchical two-pass alignment strategy implemented in 3dvolreg. First, volumes within each run were aligned to a run-specific reference image; subsequently, these reference volumes were aligned to a common reference, bringing all runs into a shared alignment space. Alignment was performed in two stages, with an initial low-resolution pass estimating gross head motion followed by refinement at full resolution. Unless otherwise specified, subsequent preprocessing steps were applied to the *backtothefuture* task.

Each volume was aligned to a reference image selected based on the minimum outlier fraction across runs. The motion-corrected functional images were then aligned to the anatomical image using align_epi_anat.py and subsequently nonlinearly transformed to MNI standard space.

A whole-brain mask was generated by computing the union of run-specific masks. Functional images were spatially smoothed to a target smoothness of 4 mm full-width at half-maximum (FWHM) using 3dBlurToFWHM to improve signal-to-noise ratio. Temporal band-pass filtering (0.01–1 Hz) was then applied to attenuate low-frequency drift and highfrequency noise.

Because the runs in the *backtothefuture* task were long, the baseline polynomial degree was fixed at two.

### Physiological

Raw pulse oximetry recordings (see Physiological recordings) were converted from the scanner’s native log format to BIDS-compliant format by extracting the cardiac waveform and trigger events from each run, computing the sampling frequency (200 Hz) from the log file metadata, and aligning the physiological time series to the fMRI acquisition using scanner timestamps.

RETROICOR-based cardiac noise regressors were then generated using AFNI’s physio_calc.py, which implements the method described by Glover et al., 2000. For each run, the program identified cardiac peaks in the pulse oximetry waveform, estimated the cardiac phase at each slice acquisition, and produced a set of Fourier-basis regressors modelling the cardiac contribution to the fMRI signal. Because the physiological recorder and MRI scanner used separate timing systems, the pulse oximetry recording occasionally ended before the final fMRI volume was acquired, resulting in regressor time series that were slightly shorter than the corresponding fMRI data (typically by 2-8 volumes; see Usage Notes for details).

The regressors were not incorporated into the primary fMRI preprocessing pipeline described above. They are provided as additional derivatives so that users may optionally include them as nuisance regressors in their own analyses.

### Data Records

Identifying information was removed from all records, and facial features were removed from anatomical images. The resulting dataset is publicly available on the OpenNeuro platform (XXX). The scripts used for data processing are available in the project’s GitHub repository (https://github.com/jbuzinel/movieproject).

#### Questionnaires

**Location** sourcedata/questionnaires.csv

**File format** Comma-separated value (CSV)

Participants’ responses to a set of questionnaires (see *Questionnaires* under *Tasks* section) were stored in a comma-separated value (CSV) file, with one row per participant and each question or test item represented as a column.

#### ESM

**Location** sourcedata/esm.csv

**File format** Comma-separated value (CSV)

Participants’ responses to ESM surveys (see *Experience Sampling Method* under *Tasks* section) were stored in a comma-separated value (CSV) file, with one row per survey and each question or test item represented as a column.

### Cognitive tasks

**Location** sourcedata/bandit_task.csv and sourcedata/selt_task.csv

**File format** Comma-separated values (CSV)

Participants’ responses to a battery of cognitive tasks (see *Cognitive Tasks* under *Tasks* section) in a CSV file. The file contains the *FID* identifier column.

#### Post-scan questionnaire

**Location** sourcedata/post_scan_questionnaire.csv

**File format** Comma-separated values (CSV)

Participants’ responses to the post-scan questionnaire (see *Post-session questionnaire* under *Data acquisition* section) in a CSV file. The file contains the *FID* identifier column.

#### Anatomical MRI

**Location** sub-<ID>/ses-<SES_ID>/anat/sub-<ID>_ses-<SES_ID>_T1w.nii.gz

**Session** 001

**File format** NIfTI, gzip-compressed

**Sequence protocol** sub-<ID>/ses-00<SES\_ID>/anat/sub-<ID>\_ses-<SES\_ID>\_T1w.json

The anatomical scan was collected as a 5-minute MPRAGE before or after completion of the movie task. The raw image had facial features removed and is provided as a 3D image file named sub-<ID>_ses-<SES_ID>_T1w.nii.gz. It is accompanied by a sequence protocol file in the same folder, saved in JSON format.

### Functional MRI

**Location** sub-<ID>/ses-<SES\_ID>/func/sub-<ID>\_ses-<SES\_ID>\_task-<TASK\_NAME>\_run-<RUN\_ID>\_bold.nii.gz

**Session** 001

**Task-name** backtothefuture

**Run** 001, 002, 003

**File format** NIfTI, gzip-compressed

**Sequence protocol** sub-<ID>/ses-<SES\_ID>/func/sub-<ID>\_ses-<SES\_ID>\_task-<TASK\_NAME>\_run-<RUN\_ID>\_bold.json

Raw fMRI data is available as individual time series files per each task run.

**Location** sub-<ID>/ses-<SES\_ID>/fmap/sub-<ID>\_ses-<SES\_ID>\_acq-<TASK\_NAME>\_ dir-PA\_run-<RUN\_ID>\_epi.nii.gz

**Session** 001

**Task-name** func

**Run** 001, 002, 003

**File format** NIfTI, gzip-compressed

**Sequence protocol** sub-<ID>/ses-<SES\_ID>/fmap/sub-<ID>\_ses-<SES\_ID>\_acq-<TASK\_NAME>\_ dir-PA\_run-<RUN\_ID>\_epi.json

The reverse phase-encoding (P→A) scan was collected for each task and is provided as a separate time series file. The task label func corresponds to the movie-watching task (referred to as *backtothefuture* in other files). The associated sequence protocol file is stored in the same directory as the functional images in JSON format.

### Freesurfer outputs

**Location** derivatives/freesurfer/sub-<ID>

Standard FreeSurfer outputs were generated by running the *recon-all* pipeline separately for each participant. The *fsaverage* directory is included alongside participant-specific files. Each participant folder also contains a *SUMA* directory produced using the *SUMA_Make_Spec_FS* command.

### SSwarper outputs

**Location** derivatives/sub-<ID>/SSwarper

Standard SSwarper outputs were generated using the AFNI *SSwarper* command applied independently to each participant. This procedure performs skull stripping of the anatomical volume and computes a nonlinear transformation to standard space.

### Preprocessed functional MRI

**Location** derivatives/sub-<ID>/<TASK_NAME>/sub-<ID>_task-<TASK_NAME>_run-<RUN_ID>_preproc.nii.gz

**Task-name** backtothefuture

**Run** 001, 002, 003

Preprocessed fMRI data were generated using the AFNI processing pipeline (*afni_proc.py*).

The data are provided as individual time-series files for each task run. The preprocessing scripts for all tasks are available in the project’s GitHub repository.

### Physiological recordings

**Location** sub-<ID>/ses-001/func/sub-<ID>\_ses-001\_task-backtothefuture\_run-<RUN\_ID>\_recording-cardiac\_physio.tsv.gz

**Session** 001

**Run** 001, 002, 003

**File format** Tab-separated values, gzip-compressed

**Sidecar**

sub-<ID>/ses-001/func/

sub-<ID>\_ses-001\_task-backtothefuture\_run-<RUN\_ID>\_recording-cardiac\_physio.json

Each file contains two columns: the cardiac waveform (arbitrary units) and a binary cardiac trigger channel (1 = trigger detected, 0 = no trigger). The JSON sidecar specifies the sampling frequency (200 Hz), the start time of the recording relative to the first fMRI volume, and column descriptions. Raw Siemens log files are retained in sourcedata.

### Physiological noise regressors

**Location**

derivatives/sub-<ID>/physio/

sub-<ID>_run-<RUN_ID>_slibase.1D

**File format** AFNI 1D text

RETROICOR cardiac noise regressors generated by AFNI’s physio_calc.py. Each file contains slice-based Fourier regressors modelling cardiac phase at each fMRI volume, suitable for inclusion as nuisance regressors in a general linear model. Quality control plots (_regressors_phys.svg) and processing logs are provided alongside each regressor file.

### Technical Validation

#### Evaluation of head motion control

To quantify in-scanner movement, framewise displacement (FD) was computed using AFNI’s 3dvolreg function. For four participants with mid-run pauses that resulted in split acquisition files (sub-07, sub-20, sub-38, sub-40), the affected runs were excluded from the head motion analysis while unaffected runs were retained. For an additional eight participants (sub-04, sub-08, sub-14, sub-22, sub-25, sub-29, sub-32, sub-33) where one or more runs were stopped early, only the volumes acquired before the interruption were included (see Usage Notes for per-participant details). Figure 2 displays the resulting distributions across all participants for each run. The data are heavily skewed towards lower values, with most observations falling below 0.3 mm. In Figure 2A, the 95th percentile, marked by a dashed red line, was 0.26 mm, confirming that the vast majority of the data reflects minimal movement. Per-run distributions, shown as violin plots in Figure 2B, were highly consistent across all three runs, with median FD below 0.1 mm and most values not exceeding 0.3 mm. Overall, these findings indicate that head motion was well-controlled throughout the scanning session, limiting the potential for motion-related artefacts in the data.

**Figure 2:**
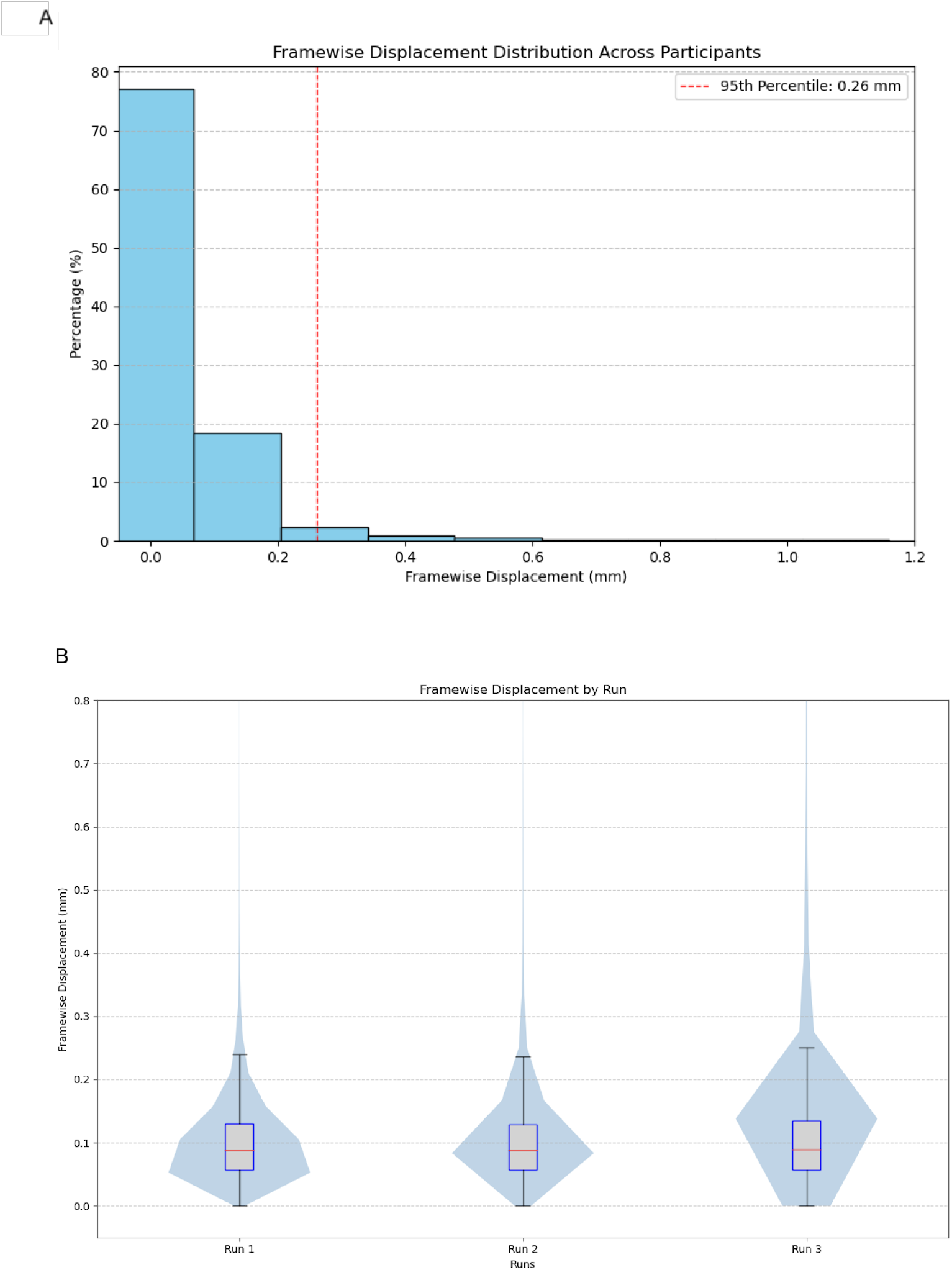
Framewise displacement (FD) distributions across all participants. The top panel (A) shows the overall FD distribution with the 95th percentile marked by a dashed red line. The bottom panel (B) shows FD distributions separately for each run, displayed as violin plots with overlaid boxplots.

Table 1 summarises head motion across the three runs of the *backtothefuture* task, reporting mean and mean maximum values for six motion parameters: three rotational (Roll, Pitch, Yaw; in degrees) and three translational (Superior-Inferior (dS), Left-Right (dL), Posterior-Anterior (dP); in millimetres). Averaged across all runs, both rotational and translational displacements stayed within ±0.09 degrees and millimetres respectively, reflecting consistently low movement throughout the scanning session. Mean maximum values remained largely below 1.0, though Pitch and dS exceeded this threshold in Runs 2 and 3, with the highest values observed in Run 3 (mean maximum Pitch = 1.67°, mean maximum dS = 1.51 mm). The same run-level exclusions described above were applied. Taken together, these results suggest that head motion was well-controlled and is unlikely to have introduced motion-related artefacts into the data.

**Table 1:** Descriptive statistics for head motion parameters across each run of the *backtothefuture* task. Mean and mean maximum values are reported for three rotational (Roll, Pitch, Yaw; in degrees) and three translational components (Superior-Inferior (dS), Left-Right (dL), Posterior-Anterior (dP); in millimetres).

| Run | Mean |  |  |  |  |  | Mean maximum |  |  |  |  |  |
| --- | --- | --- | --- | --- | --- | --- | --- | --- | --- | --- | --- | --- |
|  | Roll | Pitch | Yaw | dS | dL | dP | Roll | Pitch | Yaw | dS | dL | dP |
| 1 | -0.00 | -0.07 | -0.04 | -0.01 | -0.01 | -0.03 | 0.34 | 0.85 | 0.39 | 0.84 | 0.24 | 0.58 |
| 2 | -0.02 | -0.09 | 0.02 | -0.01 | 0.01 | -0.03 | 0.51 | 1.06 | 0.61 | 1.05 | 0.46 | 0.59 |
| 3 | -0.03 | -0.07 | -0.00 | -0.04 | -0.01 | -0.05 | 0.56 | 1.67 | 0.66 | 1.51 | 0.43 | 0.75 |

### Inter-subject correlation during movie-watching

Inter-subject correlation (ISC) analysis was performed across all participants to evaluate temporal alignment and data quality by examining shared fluctuations in brain activity (Nastase et al., 2019). ISC is widely used to quantify synchronization of neural responses during naturalistic experimental paradigms (Hasson et al., 2004; Nastase et al., 2019). Two separate ISC analyses were conducted: one across all three runs of the movie (all-runs analysis), and one restricted to run 3 only. Eleven participants were excluded from the all-runs analysis, and two participants were excluded from the run 3 analysis, due to mismatched timeseries lengths (see Usage Notes for details).

ISC values were computed using AFNI tools (*3dTcorrelate* and *3dISC*). Pairwise correlations were calculated for all unique participant pairs using Pearson correlation coefficients followed by Fisher *z*-transformation. Group-level statistical inference was conducted using a mixed-effects model in which both members of each pair were treated as random intercepts. Statistical significance was determined by correcting *t*-statistics for multiple comparisons using the False Discovery Rate (FDR) procedure with a threshold of *q <* 0.001. The resulting ISC maps are presented in Figures 3 and 4.

**Figure 3:**
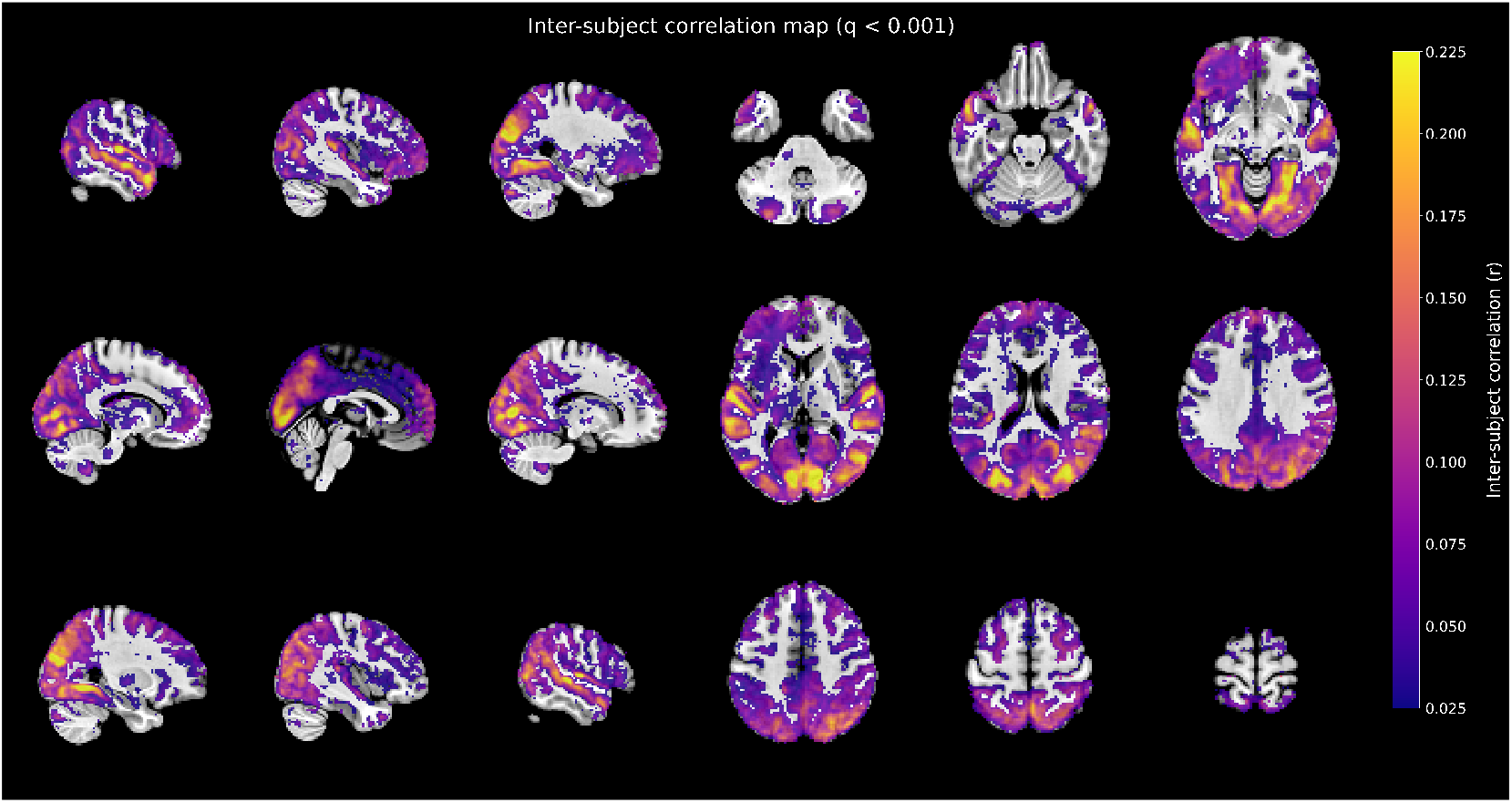
Voxelwise inter-subject correlation while viewing *Back to the Future* (all runs), calculated from 30 participants after excluding 11 due to mismatched timeseries lengths (see Usage Notes). Statistical significance was assessed with voxelwise t-tests and controlled for multiple comparisons using the False Discovery Rate (FDR) procedure at q < 0.001 (two-tailed). No additional cluster-size threshold was applied. The colour scale reflects Pearson r.

**Figure 4:**
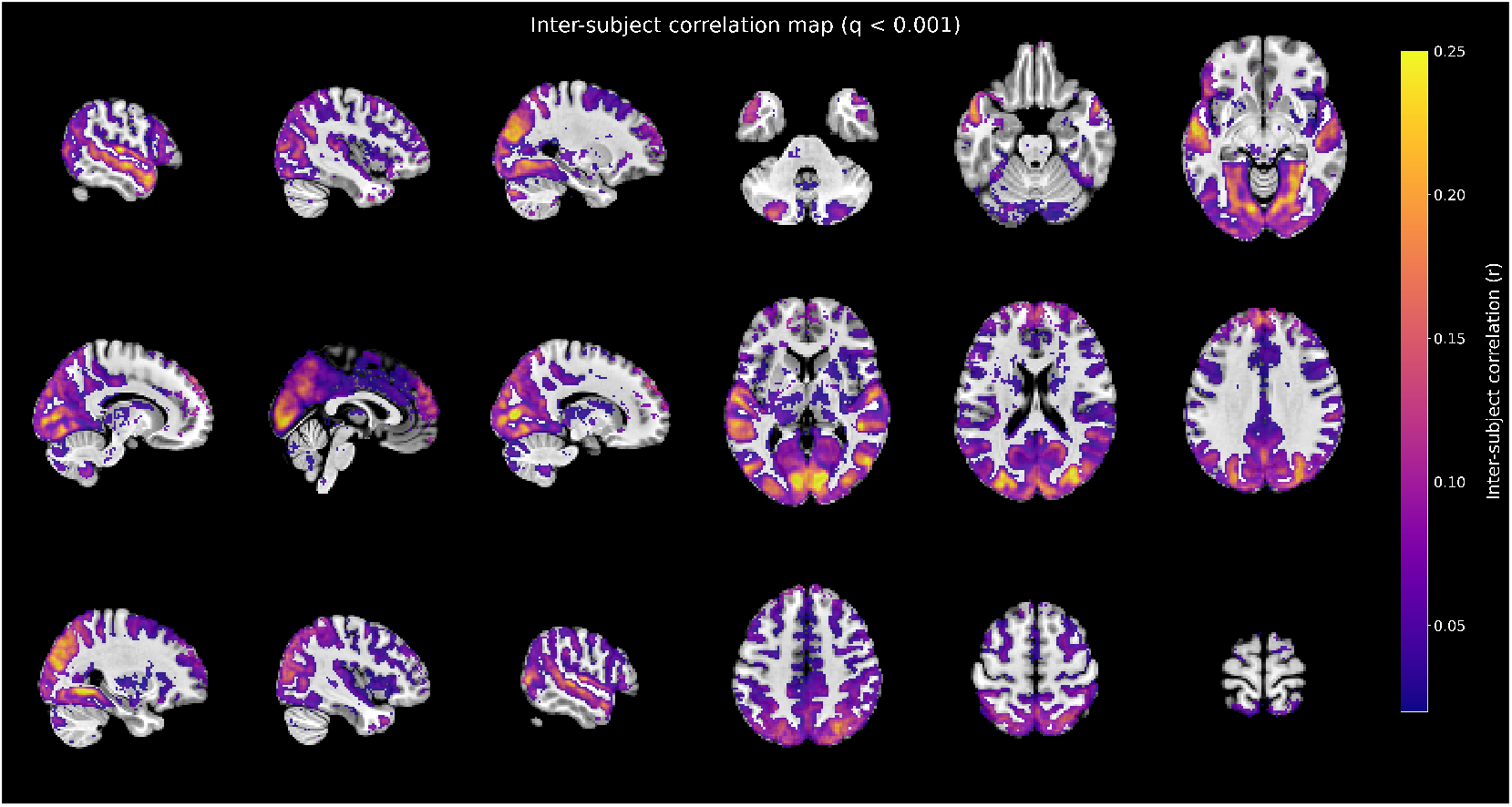
Voxelwise inter-subject correlation during *Back to the Future* viewing (run 3 only), computed across 39 participants after excluding two with run 3 data issues (sub-04: truncated run; sub-40: paused mid-scan). Statistical significance was assessed with voxelwise t-tests and controlled for multiple comparisons using the False Discovery Rate (FDR) procedure at q < 0.001 (two-tailed). No additional cluster-size threshold was applied.

Robust ISC values across early sensory cortices indicated that both stimulus presentation and preprocessing successfully preserved temporal alignment in the data. The observed ISC maps showed widespread synchronization across participants in cortical regions associated with sensory and higher-order processing, particularly the primary visual and auditory cortices, as well as posterior medial regions such as the precuneus and posterior cingulate cortex. These findings are consistent with previous naturalistic fMRI studies and likely reflect shared neural engagement with the movie stimulus (Aliko et al., 2020; Hasson et al., 2010; Lerner et al., 2011).

### Eye-tracker quality control

Participants completed a standard nine-point calibration and validation procedure prior to every run of the *backtothefuture* task. Calibration quality was assessed by extracting the mean and maximum error values for each eye and run from the corresponding EyeLink .asc files across all participants. A summary table reporting average and maximum errors for each participant, run, and eye is provided in the Supplementary Table S2. Cases in which calibration data were unavailable are reported in Supplementary Table S3.

Calibration quality was evaluated according to SR Research guidelines, where calibration was rated as *GOOD* if average error stayed under 1.0*^◦^* and the maximum error under 1.5*^◦^*, *FAIR* if average error fell between 1.0*^◦^* and 1.5*^◦^* or the maximum error between 1.5*^◦^*and 2.0*^◦^*, and *POOR* when the average error exceeded 1.5*^◦^* or the maximum error exceeded 2.0*^◦^*(https://www.sr-research.com/support/showthread.php?tid=244).

Across all calibration instances (*N* = 150), the mean average error was 0.61*^◦^* (SD = 0.41*^◦^*), while the mean maximum error was 1.43*^◦^* (SD = 1.67*^◦^*). Based on these criteria, 120 calibrations (80.0%) were classified as *GOOD*, 8 (5.3%) as *FAIR*, and 22 (14.7%) as *POOR*.

Because data were acquired using a binocular setup, run-level quality was defined such that a run was considered *GOOD* if at least one eye met the GOOD calibration criterion, *FAIR* if the best available eye met only the FAIR criterion, and *POOR* otherwise. For runs in which only one eye had calibration data, quality was determined from that eye alone. This analysis was restricted to runs where calibration could be assessed, excluding the 10 participants for whom no calibration data were recorded in any run (sub-01, sub-02, sub-03, sub-06, sub-09, sub-11, sub-12, sub-13, sub-14, sub-15; see Supplementary Table S3). Under this definition, 68 runs were classified as *GOOD*, 2 as *FAIR*, and 11 as *POOR*, indicating generally high calibration accuracy across the eye-tracking recordings.

### Usage notes

In a few cases, the data acquisition was atypical with fMRI recording being paused or interrupted during the scan. The details are outlined below:

- sub-04: run 2 was paused several times and temporal alignment cannot be guaranteed, thus the run was excluded from processing (raw data is, however, available). Due to technical reasons, run 3 was stopped 31TRs earlier. Run 1 was collected with no issues.
- sub-07: run 1 was paused (at 1088 TRs) and resumed using the built-in functionality, but due to problems with the protocol is missing the last 16 TRs. Run 2 was paused twice, with the final chunk being too short for preprocessing, meaning that run 2 is missing the final 30 TRs. Both runs are not recommended for time-sensitive analysis, although they may be used with additional checks. Run 3 was collected with no issues.
- sub-08: run 1 was paused (at 202 TRs) and resumed but due to technical reasons, stimulus alignment cannot be guaranteed after TR 202 in this run. Run 2 was interrupted and finished at TR 1081. Run 3 was collected without issues.
- sub-14: run 1 was stopped early (at 722 TRs). Runs 2 and 3 were collected without issues.
- sub-20: run 2 was paused with 833 TRs and then resumed. Due to technical issues, the run is not recommended for time-sensitive analyses. Runs 1 and 3 were collected without issues.
- sub-22: run 1 was stopped early (at 1248 TRs). Runs 2 and 3 were collected without issues.
- sub-25: run 2 was stopped early (at 1488 TRs). Runs 1 and 3 were collected without issues.
- sub-29: run 2 was paused (at 750 TRs), could not be restarted due to technical reasons, and was therefore stopped early. Runs 1 and 3 were collected without issues.
- sub-32: run 1 was stopped early (at 300 TRs). Runs 2 and 3 were collected without issues.
- sub-33: run 1 was stopped early (at 1014 TRs) due to participant discomfort. Runs 2 and 3 were collected without issues.
- sub-38: run 2 was paused (at 619 TRs) and resumed. Runs 1 and 3 were collected without issues.
- sub-40: run 3 was paused (at 1474 TRs) and resumed. Runs 1 and 2 were collected without issues.

The MATLAB script for presenting the movie stimulus included a pause function in case participants squeezed the alert ball due to any discomfort, technical (e.g., audio) issues, a desire for a break, or any other reason during the run. When the movie was paused (by pressing ‘P’ on the keyboard), it stopped on the last frame until ‘R’ was pressed to resume. Upon resuming, the movie rewound 12 seconds and paused for eight spare TRs from the scanner. The scanner operator adjusted the scanning protocol to account for the pause. In cases where the pause functionality was used, two files for relevant run were created (’*beforepause*’ and ‘*afterpause*’), and when the pause functionality was used more than once in a run, they were labelled as ‘parts’ (e.g., ‘*part1*’, ‘*part2*’, ‘*part3*’).

The database includes raw and preprocessed MRI data, and preprocessing pipelines using AFNI, Freesurfer, MATLAB and Python were designed to follow best practice for functional neuroimaging. The pipelines have been tailored to the specific demands of the current task, however, different approaches may be best suited for different hypotheses and analytical approaches. Depending on the investigation in question, users may wish to modify parts of the preprocessing such as motion censoring thresholds, spatial smoothing, filtering, or alignment methods. We encourage users to examine the GitHub repository (see *Code availability*), which provides all preprocessing scripts, and quality control reports. Reviewing these may aid with reproducibility and processing decisions, and help users create custom workflows.

In addition to the fMRI acquisition issues documented above, the following physiological recording anomalies were also identified:

- **No physiological data:** Pulse oximetry recordings were not available for sub-36, sub-39, and sub-41 across all three runs. In addition, missing pulse oximetry data were identified for the following individual runs: sub-04 run-003, sub-07 runs-002 and -003, sub-15 runs-001 and -002, sub-30 run-003, and sub-35 run-001.
- **Truncated physiological recordings:** In five cases, the pulse oximetry recording ended before the fMRI acquisition, despite the fMRI data being collected without issues for the corresponding run:

– sub-01 run-002: recording duration approximately 2163s versus 2283s expected
– sub-12 run-003: recording duration approximately 68s versus 2412s expected
– sub-13 run-003: recording duration approximately 1293s versus 2412 s expected
– sub-17 run-003: recording duration approximately 181s versus 2412s expected
– sub-30 run-002: recording duration approximately 1210s versus 2283s expected
- **Truncated physiological recordings (consistent with fMRI interruptions):** In five additional cases, the physiological recording ended early in line with documented fMRI acquisition interruptions (see above):

– sub-14 run-001: recording duration approximately 1107s versus 2040 s expected (fMRI stopped at 722 TRs)
– sub-22 run-001: recording duration approximately 1893s versus 2040s expected (fMRI stopped at 1248 TRs)
– sub-29 run-002: recording duration approximately 67s versus 2283s expected (fMRI paused at 750 TRs and stopped)
– sub-32 run-001: recording duration approximately 472s versus 2040s expected (fMRI stopped at 300 TRs)
– sub-33 run-001: recording duration approximately 1543s versus 2040s expected (fMRI stopped at 1014 TRs)

RETROICOR regressors were not generated for these runs. The raw (truncated) physiological recordings are included in the dataset for completeness.

- **Split physiological recordings:** For participants with mid-scan pauses (sub-04, sub07, sub-08, sub-20, sub-38, sub-40), the physiological recordings are split into multiple files per run (labelled beforepause/afterpause or part1/part2/part3 in sourcedata). These segments correspond to the fMRI acquisition pauses documented above.
- **Regressor coverage:** For runs where RETROICOR regressors were successfully generated, the regressor time series may contain slightly fewer time points than the corresponding fMRI data (typically 2–8 fewer volumes). This was due to the pulse oximetry recording stopping a few seconds before the final fMRI volume was acquired. Users incorporating these regressors should ensure that the fMRI and regressor time series lengths are matched, for example by zero-padding the regressors.

### Limitations and Future Directions

The movie stimulus could not be included in the dataset due to copyright restrictions. The movie can, however, be purchased with the International/European Article Number (EAN: 5050582401288) or the unique Amazon Standard Identification Number (ASIN: B000BVK82I). Researchers may wish to apply time-coded annotations using publicly available tools to a legally obtained copy of the movie. When doing so, the frame-level metadata and scanner trigger messages provided in the NNDb-Teens dataset can be used. The movie stimulus used in this dataset is exactly the same as the one used in NNDb v1.0 and NNDb-3T+ datasets, and all extracted annotations from the previous versions are entirely applicable to the current dataset.

Recent advances in machine learning may provide ample opportunities for investigating different aspects of the movie, such as dialogue, visual objects, faces, and emotional tone. This can be achieved through tools like Neuroscout (https://neuroscout.org), which provide frame-by-frame annotation for various popular movies, including *Back to the Future*. Such tools can also be adapted for the goals and requirements of individual projects. Importantly, however, analyses of the dataset do not need to rely on annotations. Several approaches to naturalistic neuroimaging can be conducted without predefined annotations, including inter-subject functional connectivity (Simony et al., 2016), inter-subject representational similarity (Nastase et al., 2019), dynamic functional connectivity (Allen et al., 2014), and model-free analyses of temporal reliability (Hasson et al., 2010). Thus, the dataset can support a range of analyses beyond annotation-based approaches.

Another limitation of the dataset is that it uses a single movie, which restricts the range of scenes, events, and semantic contexts available for analysis. This may reduce statistical power for some analyses and limit the generalisability of findings beyond the particular narrative and properties of the selected movie. Studies targeting low-frequency events, such as non-verbal social exchanges, ambiguous language, or non-human sounds, may therefore be underpowered.

Last, the participant sample also carries limitations worth noting. The dataset consists of 41 adolescents, which, while sufficient for many analyses, constrains the examination of individual differences or subgroup effects. The sample is predominantly female (n = 31, 75.6%), which may also limit the generalisability of findings across sexes, while the inclusion of both left- and right-handed participants should be considered when investigating lateralised processes. On average, participants reported mild symptoms of depression as measured by the PHQ-8 (mean(SD) = 9.12(5.21)) and anxiety as measured by the GAD-7 (mean(SD) = 7.54(4.66)). Additionally, a proportion of participants may suffer from lower-quality data due to pausing artifacts during certain runs and the related uncertainty regarding stimulus alignment, which may introduce noise or require data exclusion depending on the analytic approach. That said, the vast majority of participants and runs are of high quality, meaning that these issues are not expected to significantly undermine the dataset’s overall utility.

## Conclusion

In summary, the NNDb-Teens dataset provides a rich, openly available resource for studying the adolescent brain under naturalistic conditions. To our knowledge, it is the first publicly available adolescent dataset to combine full-length movie-watching fMRI with concurrent eye-tracking and physiological monitoring alongside extensive behavioural and cognitive assessments. Technical validation confirms high data quality, with low head motion, strong inter-subject neural synchronisation, and generally good eye-tracker calibration across participants. The BIDS-compliant format and publicly available processing code further support transparency and reproducibility. We anticipate that this dataset will facilitate a wide range of investigations into adolescent brain function, from sensory and social processing to individual differences in mental health and cognition. When used alongside the adult NNDb-3T+ dataset, it also opens the door to cross-sectional comparisons of naturalistic neural processing across development. We hope that by making these data freely accessible, we can support and accelerate collaborative, reproducible research into the developing brain during this critical period.

## Code availability

All the code is available on the GitHub repository: https://github.com/jbuzinel/movieproject/.

## Acknowledgements

This work was supported by the Prudence Trust and the Birkbeck-UCL Centre for NeuroImaging (BUCNI). All MRI scanning hours were donated by BUCNI as part of its commitment to open science resources. We would like to thank Letitia Schneider, Roger Atkins, Raha Razin, Oliver Josephs, Joerg Magerkurth, Winnie Yeh and other staff at BUCNI for their support with scanning, Yumeya Yamamori for his support with behavioural task development, Amy Polglase, Adam Talib, Emily De Groot and Katy Whadcock for support with data collection, and the participants and their families for their time and participation in this study.

## Supplementary materials

**Table S1.** Data availability for each participant.‘+’ indicates that the participant fully completed the assessment; ‘-’ indicates that the data is either missing or incomplete. All participants have a PHQ-8 and GAD-7 questionnaire score; those with ‘-’ under Questionnaires are missing at least some of the remaining questionnaires from the battery.

| Sub-ID | Questionnaires | ESM | Cognitive tasks | Movie-watching |
| --- | --- | --- | --- | --- |
| 1 | - | - | - | + |
| 2 | - | - | - | + |
| 3 | - | - | - | + |
| 4 | + | - | + | + |
| 5 | - | - | - | + |
| 6 | + | + | + | + |
| 7 | + | + | + | + |
| 8 | + | + | + | + |
| 9 | + | + | + | + |
| 10 | + | + | + | + |
| 11 | + | + | + | + |
| 12 | + | + | + | + |
| 13 | - | + | + | + |
| 14 | + | + | - | + |
| 15 | + | + | + | + |
| 16 | + | + | + | + |
| 17 | + | + | + | + |
| 18 | + | + | + | + |
| 19 | + | + | + | + |
| 20 | + | + | + | + |
| 21 | + | + | + | + |
| 22 | + | + | + | + |
| 23 | + | + | + | + |
| 24 | + | + | + | + |
| 25 | + | + | + | + |
| 26 | + | + | + | + |
| 27 | + | + | + | + |
| 28 | + | + | + | + |
| 29 | + | + | + | + |
| 30 | + | + | + | + |
| 31 | + | + | + | + |
| 32 | + | - | + | + |
| 33 | + | - | + | + |
| 34 | + | - | + | + |
| 35 | + | - | + | + |
| 36 | + | + | + | + |
| 37 | + | + | + | + |
| 38 | + | + | + | + |
| 39 | + | - | + | + |
| 40 | + | - | + | + |
| 41 | + | + | + | + |
| **Total N** | **36** | **30** | **36** | **41** |

**Table S2.**
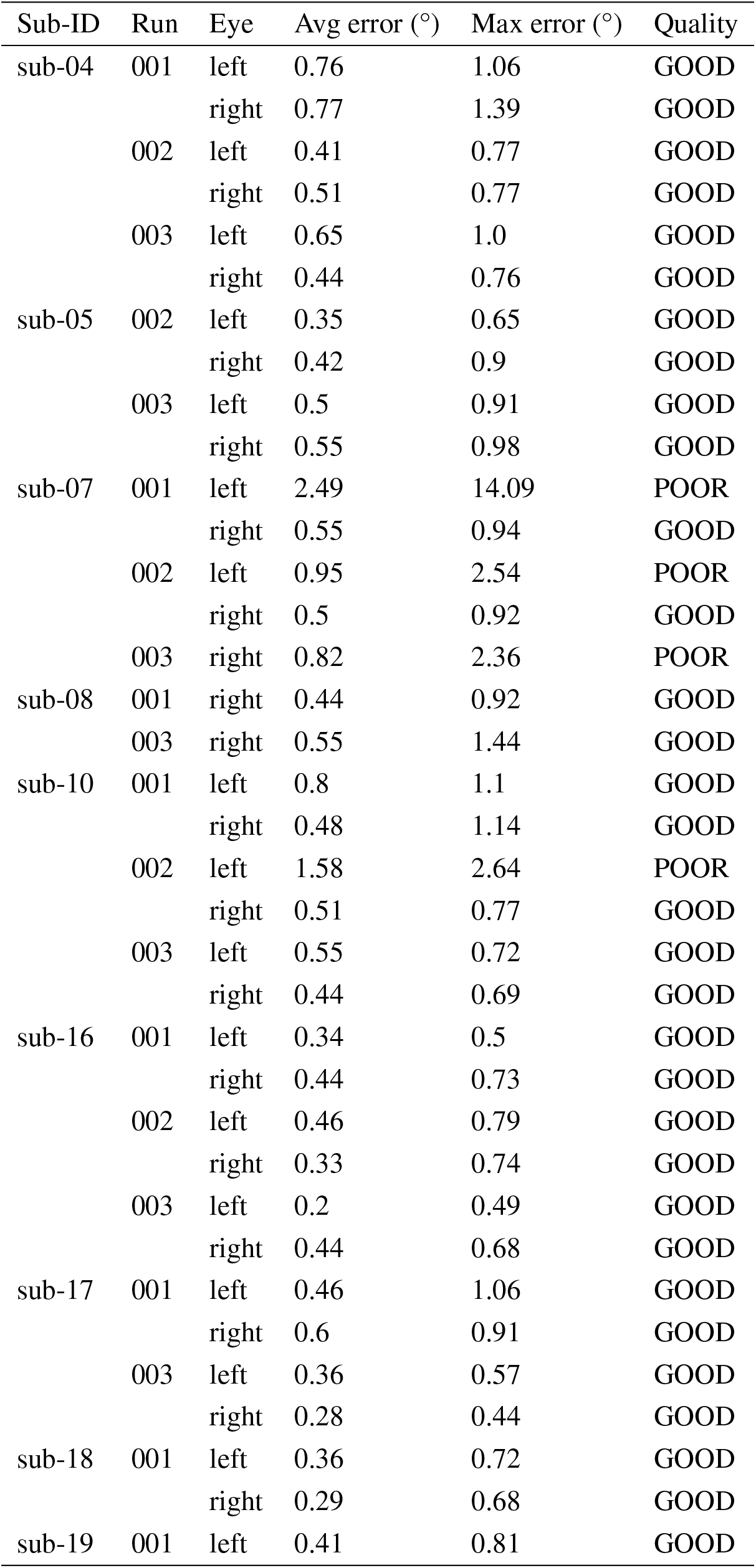
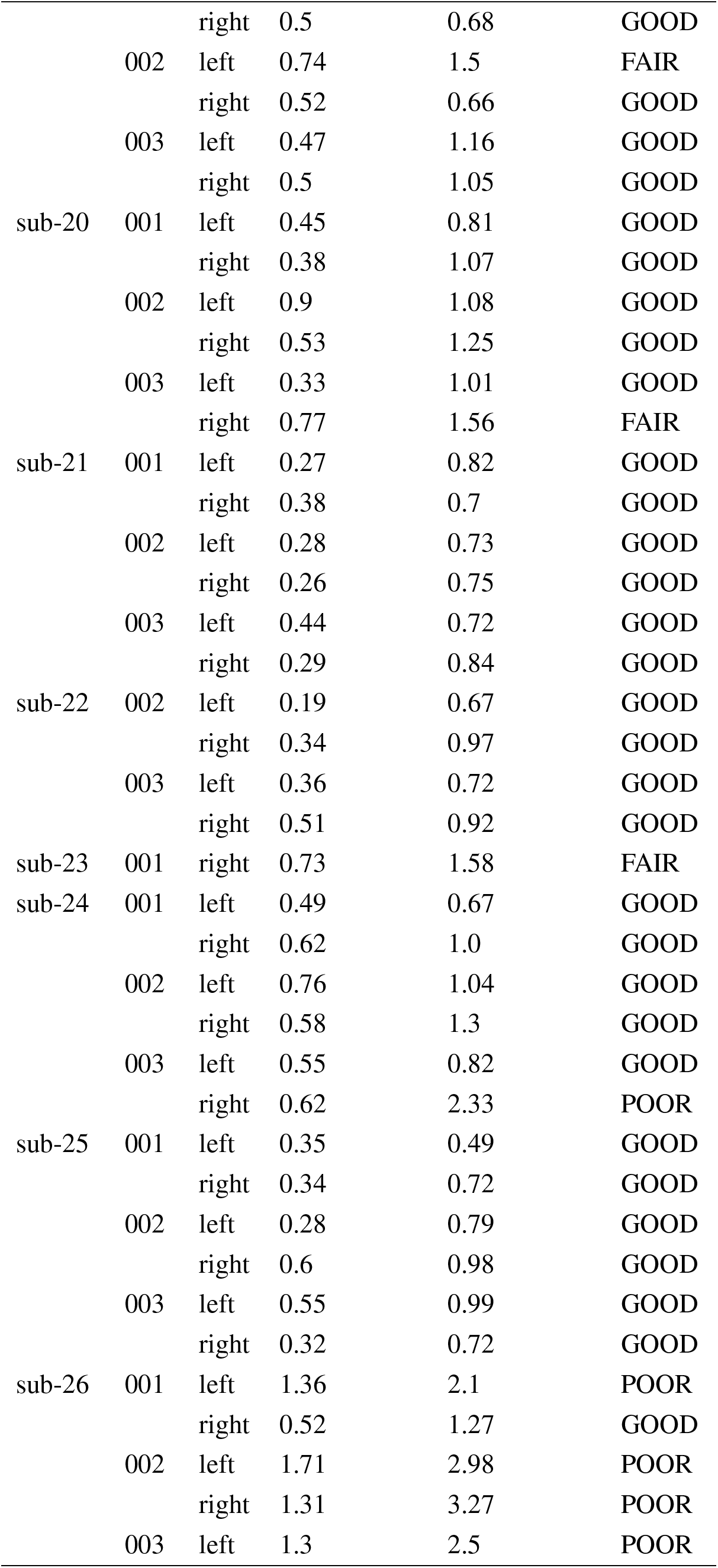
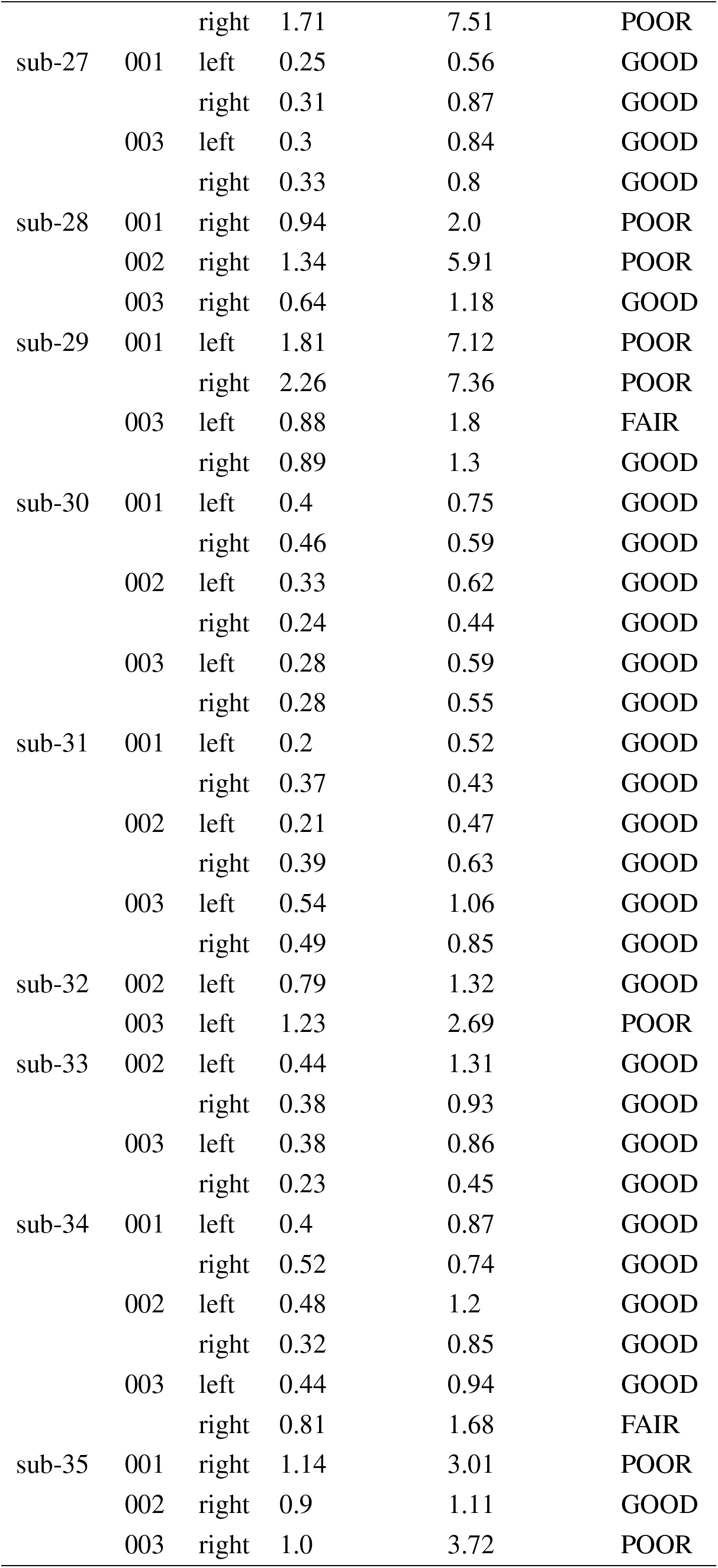
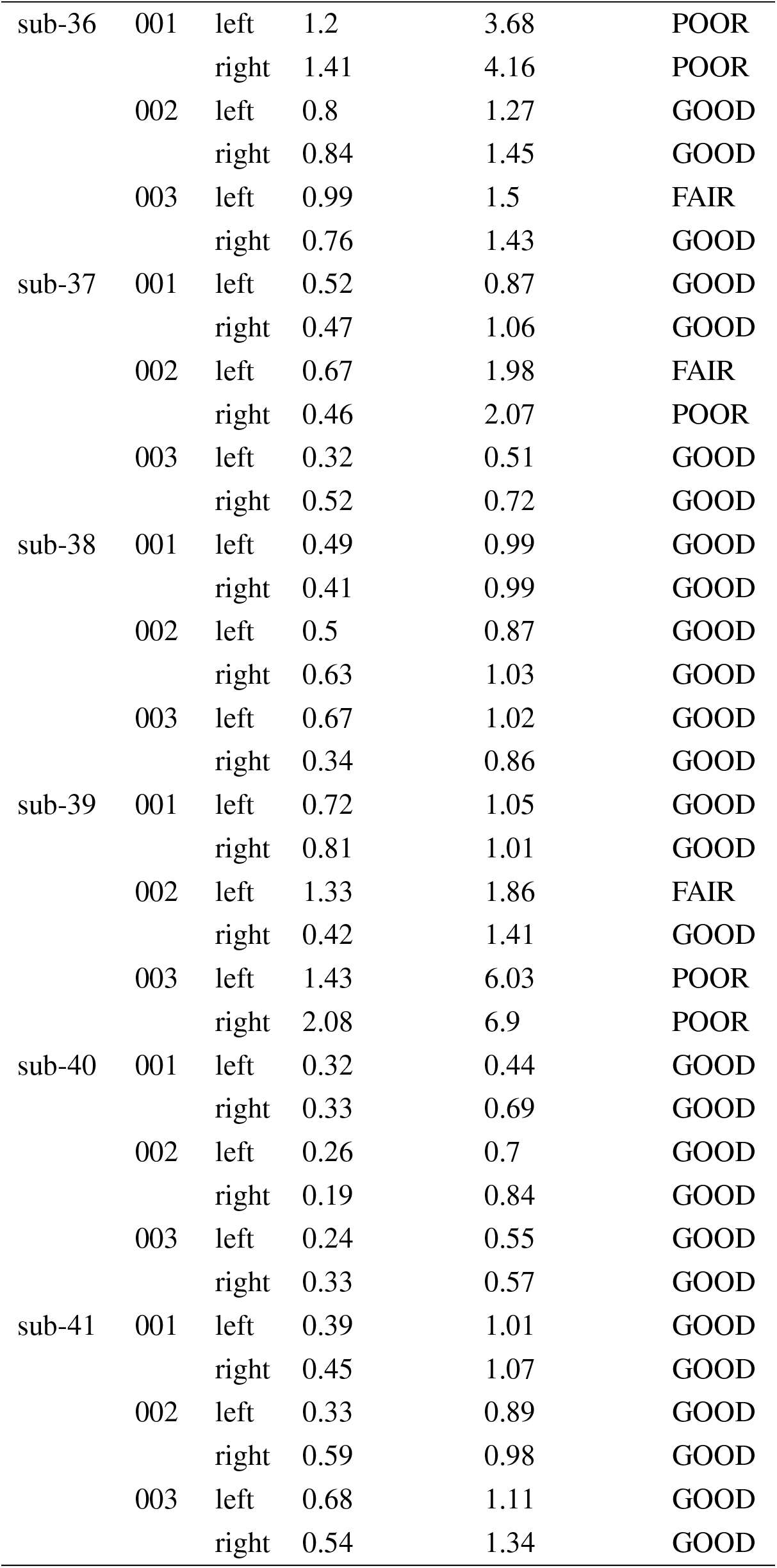
Eye-tracker calibration quality for each participant, eye, and run of the *backtothefuture* task. Quality ratings follow SR Research guidelines: GOOD = average error < 1.0° and maximum error < 1.5°; FAIR = average error 1.0–1.5° or maximum error 1.5–2.0°; POOR = average error *>* 1.5° or maximum error *>* 2.0°.

**Table S3.**
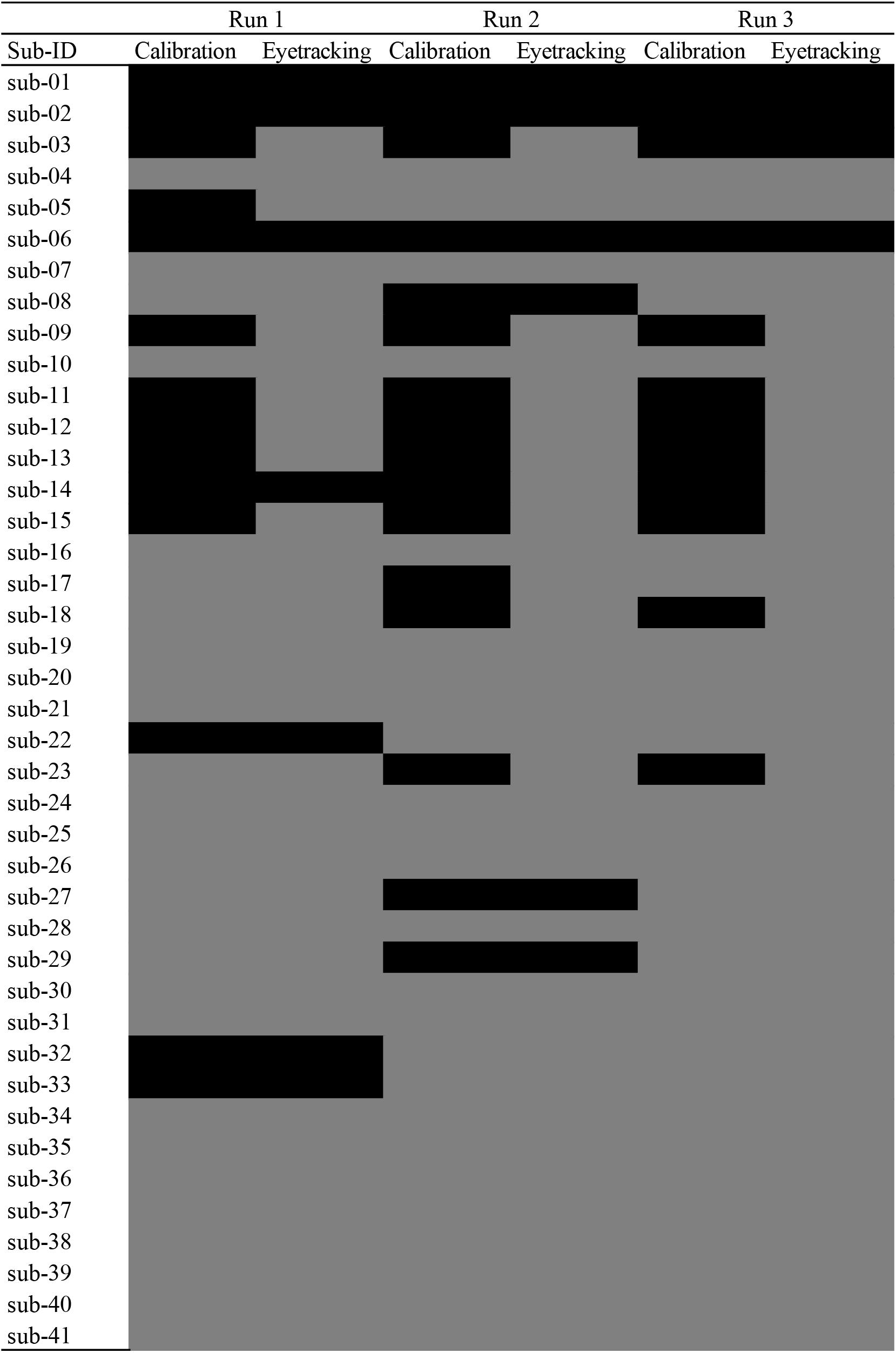
Availability of eye-tracking data. Grey cells indicate that data are available; black cells indicate that data are missing. Where a calibration file is available from a previous run for a given participant, it can be applied to subsequent runs.

